# FECAL MICROBIOME OF CHILDREN WITH NOROVIRUS GASTROENTERITIS

**DOI:** 10.1101/2021.03.23.436635

**Authors:** Juliana Merces Hernandez, Edivaldo Costa Sousa Junior, Giovanna Brunetta Sant’Ana Almeida, Ana Caroline Rodrigues Portela, Maria Silvia Sousa Lucena, Jedson Ferreira Cardoso, Tammy Kathlyn Amaral Reymão, Clayton Pereira Silva Lima, Márcio Roberto Teixeira Nunes, Dielle Monteiro Teixeira, Jones Anderson Monteiro Siqueira, Yvone Benchimol Gabbay, Luciana Damascena Silva

## Abstract

The human fecal microbiome is composed of endogenous bacteria, eukaryotic viruses, bacteriophages and retroviruses. Several pathological conditions, including gastroenteritis, may be characterized by imbalance of gastrointestinal functions, with alteration in the diversity and composition of the fecal microbiota. Were analyzed twenty-seven fecal microbiome in children hospitalized with gastroenteritis (norovirus positive) from northern region of Brazil. After sequencing, was verified the presence of the domains Bacteria (95%) and Eukaryota (3.1%), the viruses represented 1.9%. Among the pathogenic viruses were found in addition to noroviruses the picornaviruses, enterovirus and parechovirus. The bacteriophages detected were of Caudovirales order, families *Siphoviridae*, *Podoviridae* and *Myoviridae*. In 22.2% (6/27) of the samples was observed co-infection between norovirus, enterovirus B and echovirus. As for the others components of the microbiome, we can highlight the presence of the taxonomic groups: Terrabacteria (50.2%), composed mainly of Actinobacteria and Firmicutes; Proteobacteria (34.5%) represented by the *Enterobacteriaceae* family; and FCB group (22%) whose most abundant microorganisms were those of the phylum Bacterioidetes. We performed a metagenomic approach to analyze the fecal microbiota of children with viral gastroenteritis, it was observed that the bacterias (*Enterobacteriaceae*) deserve attention in a possible association with noroviruses, as they were found in large quantities in infections. In addition, other enteric viruses were observed, such as enteroviruses.

## INTRODUCTION

The fecal microbiota is a complex community formed by diverse biological components that reflects the gut microbiota, a critical component of human health which contributes to immunity, nutrition, and behavior (Thursby & Juge 2017). The human gut microbiota is composed by a complex and dynamic population of microorganisms, such as bacteria, archaea, eukaryotic microbes, and viruses, which plays vital functions on the host during homeostasis and disease (Carding et al., 2017).

Studies have shown that the human intestinal microbiota is composed for four main phyla that characterize the dominant human intestinal microbiota: Firmicutes, Bacteroidetes, Actinobacteria and Proteobacteria. The gut microbiome in a health person form a balance ecosystem with important physiological functions like metabolic functions and protect against invading microorganisms (Shreiner et al., 2015). The disruption of this homeostatic system may be promoted by infectious agents. Thereby, the gastrointestinal infections can cause dysbiosis (microbial imbalance) which can affect disease prognosis (Jackson et al., 2018).

The metagenomic approach facilitated the knowledge of the microbiota including the virome of an infection that was largely unknown due to the difficulty in cell culture of these infectious agents (Riesenfeld et al., 2004). The virome of gut microbiome is composed basically by phages, mainly of the order *Caudovirales.* Within this order, the families *Myoviridae*, *Siphoviridae* and *Podoviridae* are the most frequent (Hoyles et al., 2014; Castro-Mejía et al., 2015).

Diarrheal diseases are the second leading cause of death for children under five, causing 525,000 children each year (WHO, 2020). Human noroviruses are a major cause of viral gastroenteritis and are the main etiological agents of acute gastroenteritis (AGE) outbreaks. This disease is related to intestinal inflammation and it is estimated that norovirus are associated to around 200,000 deaths per year, accounting for more than 90% of gastroenteritis outbreaks worldwide (Bányai et al., 2018). The cases are most often reported in health care settings and people of all ages can be affected, but children, elderly and immunocompromised people can have a severe disease (Robilotti et al., 2015).

Norovirus belongs to the family *Caliciviridae*, are a genetically diverse viruses that have positive-sense, single-stranded RNA genome of approximately 7.6 kb disease (Green, 2013). The genus *Norovirus* showed a wide genetic diversity, being classified into 10 genogroups (GI to GX), that are further classified into 45 genotypes (ICTV, 2020). The genogroups GI, GII and GIV infect humans, among which the GII.4 genotype has been most associated with outbreaks and sporadic cases of viral gastroenteritis (Chhabra et al., 2019).

Increasingly studies are reporting the role of human microbiota in health and disease, and it is evident that commensal bacteria play an important role in regulating and protecting human health (Davenport et al., 2017). However, few studies have examined the constitution of microorganisms present in the feces of humans with intestinal infections of viral etiology, with data of viral and bacterial populations (Yuan et al., 2020).

Thus, considering the importance of diversity studies of the human fecal microbiome, the aim of this study was to analyze fecal microbiome in children with norovirus gastroenteritis in northern Brazil. The study sought to describe the individual variations and the diversity of bacterial communities in the feces of cases of positive NoV.

## MATERIAL AND METHODS

### Clinical specimens

This study involved metagenomics approach to elucidate the microbial diversity in fecal samples from children (< 10 years old) with norovirus gastroenteritis, during a period of four years (2012 – 2016). The fecal specimens (N=27) were collected of inpatients attended in public health facilities by the National Program for Surveillance of Gastroenteritis (IEC/SVS/MS) in Belem and Manaus city, Amazon region, Brazil as described previously by Hernandez et al., 2020. The samples were previously characterized as positive for norovirus using RT-PCR and Sanger sequencing. This study was approved by the Ethics Committee on Human Research of IEC (CAAE 46841715.8.0000.0019).

### Nucleic acids extraction

In order to avoid loss of genetic material, no initial treatment was performed on the samples. A fecal suspension (10% w/v) in Tris/HCl/Ca^+2^ buffer was subjected to extraction *in house* using the silica method or QIAamp viral RNA mini kit (Qiagen) according to the manufacturer’s guidelines.

### Double-strand cDNA synthesis and NGS

The RNA was subjected to reverse transcription and second strand synthesis using the cDNA synthesis system kit (Roche). The acid nucleic quantification (RNA and cDNA) was performed by fluorimetry with Qubit 2.0 fluorometer, using the Qubit RNA BR assay kit (Thermo Fisher). The analytical evaluation of sample quality was performed by automated electrophoresis in 2100 Bioanalyzer (Agilent) as recommended by the manufacturer.

Sequencing libraries were prepared from 10-15 ng of cDNA, using the Illumina Nextera XT DNA Library Prep kit and sequenced on Illumina HiSeq 2500 instrument with the high-output V4 2×100-bp sequencing kit.

### Bioinformatic Analysis

#### Preprocessing and genome assembly

Raw sequence reads were submitted to quality check and trimming to remove the adapters using Trim_Galore tool. Low sequencing quality tails were trimmed using Phred quality score 30.

The binning was performed by two different methods, for taxonomic analysis of microbial communities using metagenomics classifiers: the reads passed directly to classification using Kraken 2 for 16S classification with Silva rRNA database (Quast et al., 2013); the reads were assembled into contigs to then perform the taxonomic classification. Taxonomic classification of every read was performed for quantitative screening of the microbial diversity obtained in the sequencing.

For taxonomic classification with contigs was used the Diamond v0.9.19.120 program, with mapping to NR database (NCBI non-redundant protein database. The genome assembly was performed with IDBA-UD algorithm. This approach is based on the de Bruijn graph, supports unequal coverage depth and was used for reconstruct longer contigs with higher accuracy (Peng et al., 2012).

The graphical plotting was performed by the Krona tool (Ondov et al., 2011), which allowed the interactive visualization of the data and estimation of microorganism abundance.

For viral genomes, the comparative analyzes were inspected to filter the contigs with viral similarity, being performed by scripts in house. Subsequently, was performed curation using the Geneious v.9.1.8 program (Kearse et al., 2012), obtaining consensus sequences of the validated viral genomes.

### Statistical methods

In the analysis, we retained the count for the most abundant bacterial phylum proportions. The proportion for each taxon was obtained with the use of in-house scripts to identify phyla, families and genus.

Along with bacterial proportions, we computed at a phylum level the bacterial diversity within samples using Shannon index, which represents the presence and the abundance of each taxa (Chiu & Chao 2016). We also correlated the presence of enteric viruses to the inter-individual variation in microbial composition and diversity.

The Shannon-Wiener Index, Spearman correlation coefficient and simple logistic regression analysis was utilized with P-values < 0.05 considered statistically significant. All statistical analyses were performed in Bioestat 5.0 (Ayres et al, 2007).

## RESULTS

In this study were assessed the microbial community composition in twenty-eight fecal specimens from children (< 10 years old) with norovirus gastroenteritis using whole genome sequencing. The deep sequencing generated approximately 317 million paired-end readings (out of 6.3 to 16.5 million readings per sample), of which about 2 million of reads were from norovirus (range from 10 to 138 thousand readings).

Contigs obtained after assembly showed the presence of two main domains, Bacteria with 95% (57540287 counts), and Eukaryota presented 3.1% (1885240) of a total of 60,564,491 contigs; viruses represented a small portion of the microbiome with only 1.9% (1138964).

For viral contigs, besides norovirus was observed the presence of other pathogenic enteric viruses, such as enterovirus B and human parechovirus (data not yet published). In 22.2% (6/27) of the samples was observed the association of norovirus and picornavirus contigs. The other components of virome are also found like minireovirus, rhabdovirus and unclassified bacteriophages (Figure 1).

**Figure 1.**
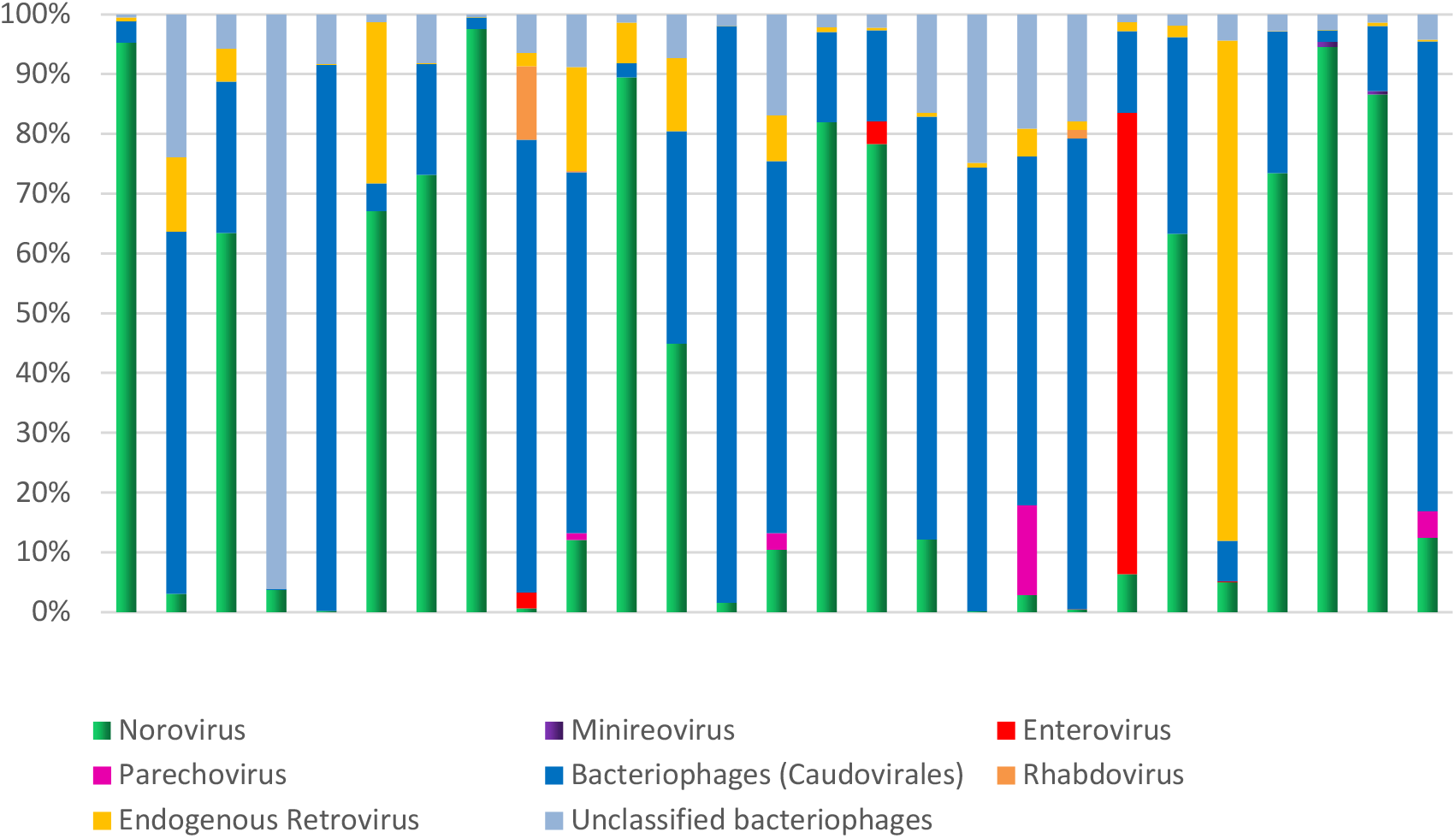
Diversity and abundance of fecal virome in children hospitalized with norovirus gastroenteritis from northern region of Brazil. Relative demonstration per sample of the components of virome.

In addition, bacteriophages and human endogenous retroviruses were also observed frequently. In the analysis of the samples individually showed the abundance of bacteriophages of the order *Caudovirales*, from the families *Siphoviridae*, *Podoviridae* and *Myoviridae*.

Among the Bacteria domain, the most abundant phylum were Firmicutes (86.3%) and Proteobacteria (12.5%). Actinobacteriota, Bacteroidota and Cyanobacteria were less representative, corresponding together to 1.2% of the mapped reads. The interindividual variations of microbial composition were estimated by shannon wiener’s index, evaluating the aspects of richness and equitability, which concern the number and proportion of each bacterial phylum in the analyzed samples.

The high interindividual variation observed reflects the community composition and can be driven by the abundance of dominant phyla (Figure 2). The data demonstrate an disequilibrium of the microbiota, mainly due to the high proportion of proteobacteria (above 5%) observed in 27% (7/26) of the analyzed samples. The statistical analysis performed between bacterial phyla showed that the proportions of Proteobacteria and Actinobacteriota were correlationed (R-Pearson= 0.5614 p=0.0028). Similarly, there was a correlation between the phyla Bacteroidota and Cyanobacteria (R-Pearson= 0.5191 p=0.0066).

**Figure 2.**
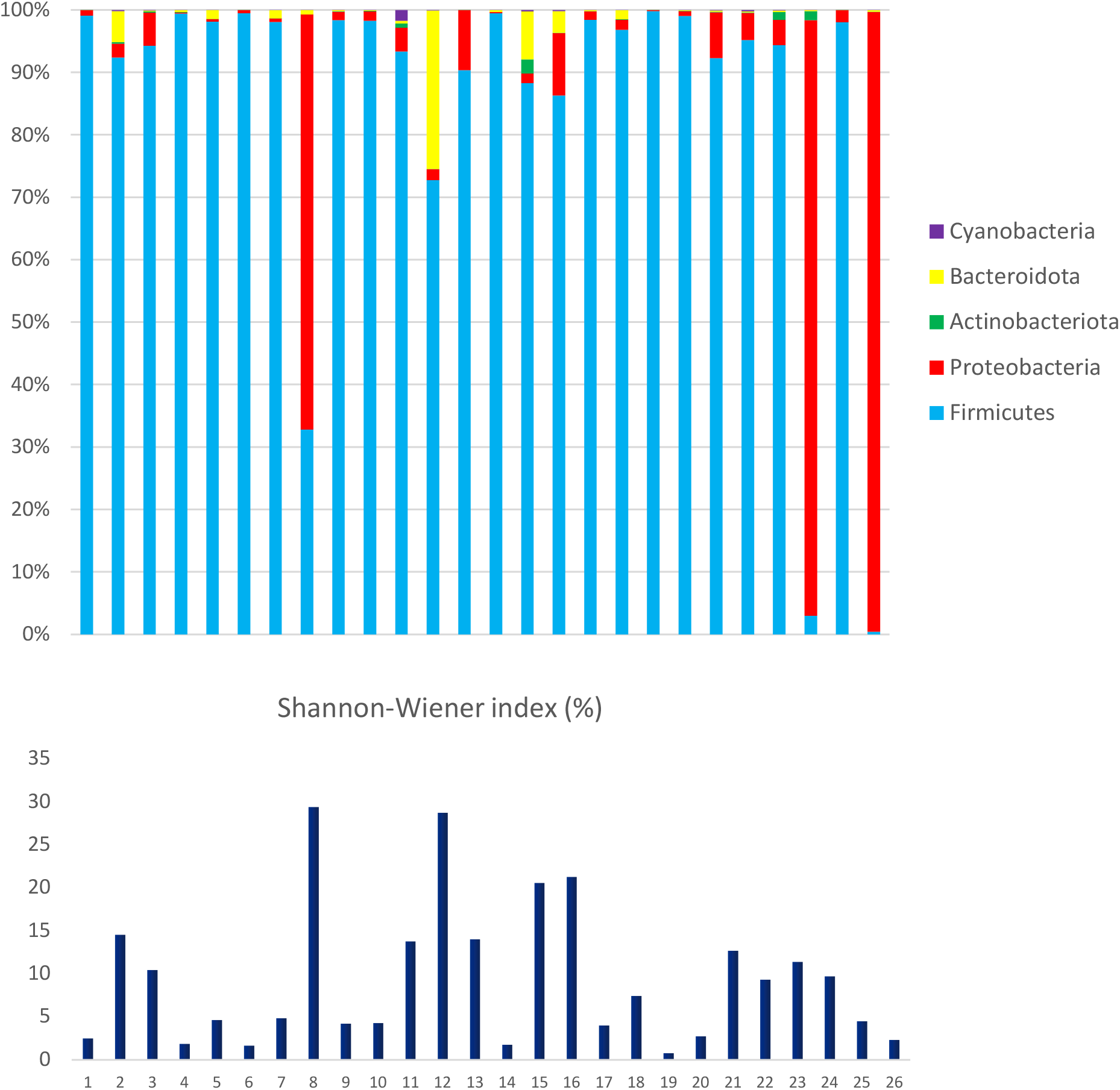
Diversity and abundance of fecal microbiome in children hospitalized with norovirus gastroenteritis from northern region of Brazil. Relative demonstration, per sample, of the main bacterial phyla that compose the human fecal microbiome and comparison with Shannon-Wiener’s index.

**Figure 3.**
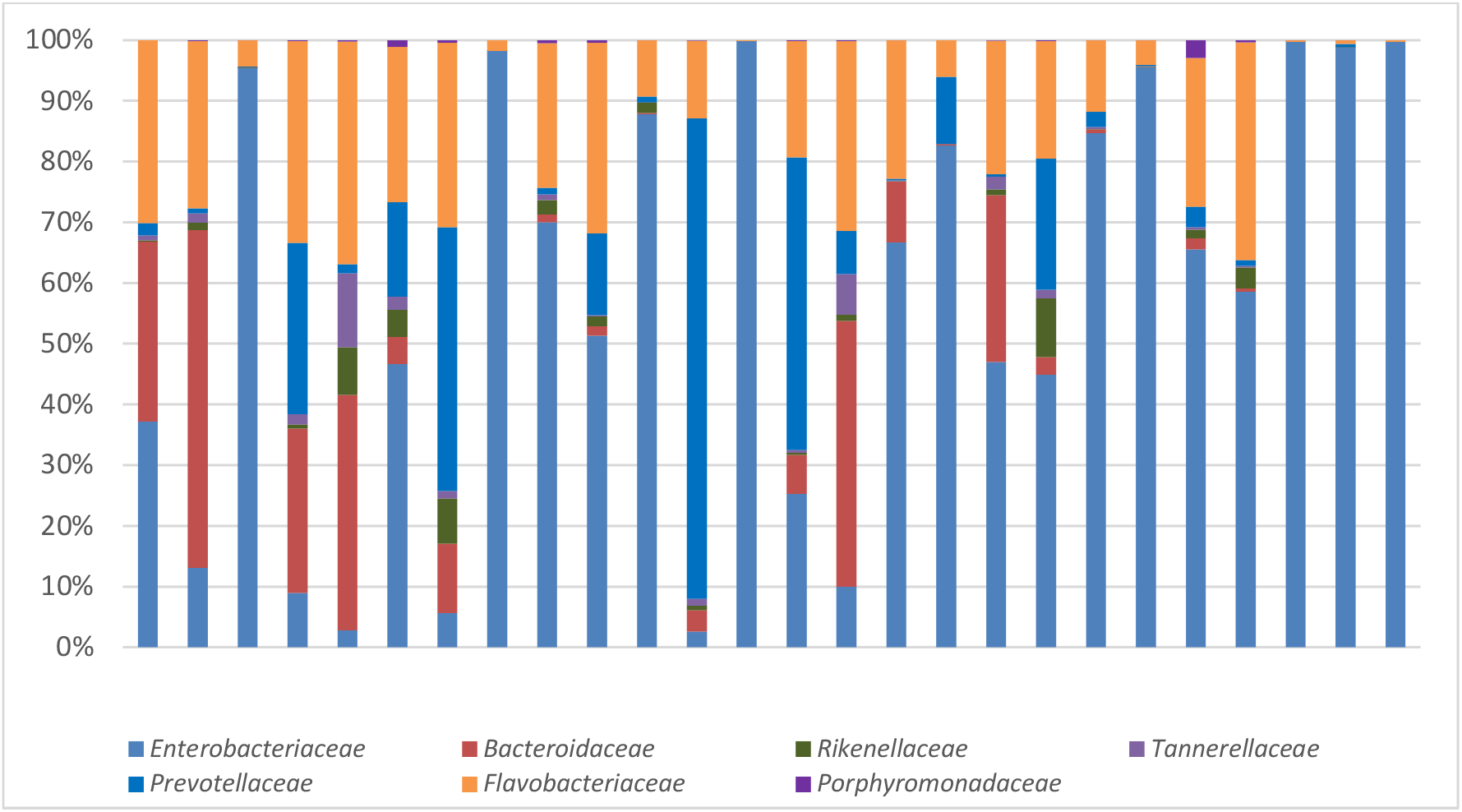
Diversity and abundance of fecal microbiome in children hospitalized with norovirus gastroenteritis from northern region of Brazil. Relative demonstration, per sample of the main bacterial families that compose the human fecal microbiome.

Considering the composition of the intestinal microbiota, a comparative study was also carried out to analyze the taxonomic profile of the analyzed samples, describing the main bacterial families detected in human fecal metagenomes. Thus, the following bacterial families were considered, which belong to the Proteobacteria and Bacteroidetes phyla: *Enterobacteriaceae*, *Bacteroidaceae*, *Rikenellaceae*, *Tannerellaceae*, *Prevotellaceae*, *Flavobacteriaceae*, *Porphyromonadaceae*.

According to the observed the *Enterobacteriaceae* family was predominant (89,5%), includes genus *Escherichia-Shigella (1.8%)*, *Klebsiella (91.4%)*, *Salmonella (0.4%)*, *Enterobacter (2.7%),* other *enterobacteria (3.7%)*. *Bacteroidaceae*, *Prevotellaceae* and *Flavobacteriaceae* was observed in 2%, 3.3% and 5 % of reads mapped; *Rikenellaceae, Tannerellaceae* and *Porphyromonadaceae* together accounted for 0.2% of reads. Therefore, the individual analysis of the samples shows the presence of possible pathogenic bacteria, the highest abundance being found for *Klebsiella* (Figure 4).

**Figure 4.**
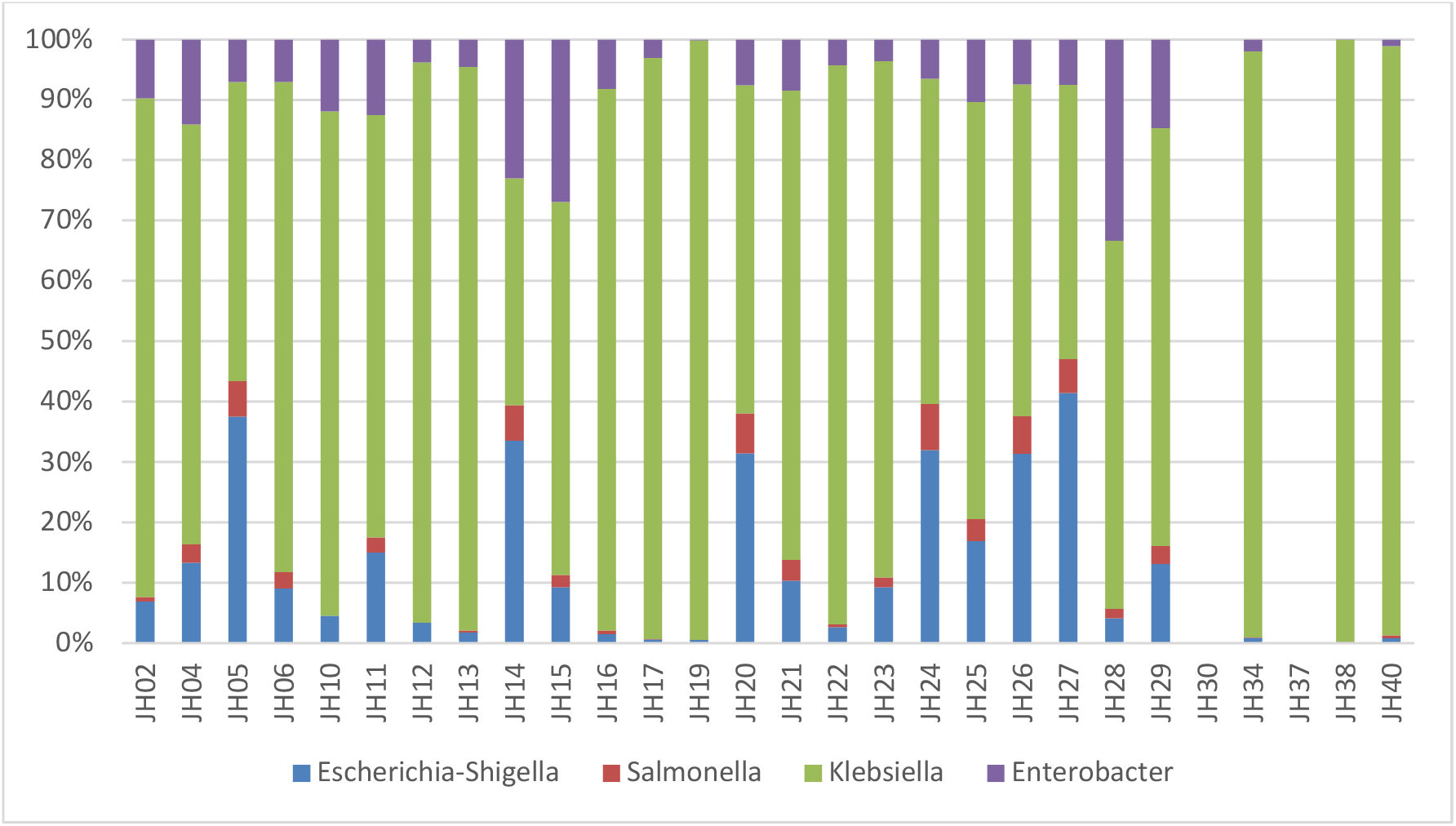
Diversity and abundance of fecal microbiome in children hospitalized with norovirus gastroenteritis from northern region of Brazil. Relative demonstration, per sample of the main genus of the family *Enterobacteriaceae*.

The direct comparison between Shannonn index of the samples with NoV contigs and samples with NoV and other viral contigs evidenced that in the samples with more than one pathogenic virus showed a less diversity, however, Mann–Whitney U test was not statistically significant (p = 0.0944).

## DISCUSSION

The human gastrointestinal tract is constituted by a larger number and variety of commensal microorganisms (bacteria, viruses, fungi, protozoa) collectively referred such as microbiome (Gigliucci et al., 2018). The fecal microbiota reflects the gut microbiome, which is composed for 90% of anaerobic bacteria in healthy adults (Bäckhed et al., 2005). These microorganisms have a mutual beneficial relationship with the host, being considered of great importance, since they act in the digestive and immunological regulation (Davenport et al., 2017).

The gastrointestinal infections change this homoeostatic balance and can lead to dysbiosis, resulting of alterations in the microbiota composition (Gigliucci et al., 2018). In the present study, we aimed to use metagenomics as a tool to investigate the composition of the intestinal microbiota in patients with an infectious disease of the gastrointestinal tract - NoV infection. Shotgun metagenomic sequencing was used to perform a taxonomic classification of the microbial communities present in fecal samples of AGE hospitalized children (0-10 years old) during a period of four years (2012 – 2016).

The children in this study showed a high abundance of Firmicutes and Proteobacteria. Were observed a high frequency of *Enterobacteriaceae* family bacteria as well as other enteric viruses such as picornavirus (Enterovirus B and Echovirus), probably due to co-infection with NoV.

The *Enterobacteriaceae* family was the most abundant and diverse pathogenic bacteria were detected with the following genus standing out, *Escherichia*, *Shigella*, *Salmonella, Klebsiella,* and *Enterobacter*. This peculiarity was also observed in other studies, which also highlighted the presence of enterobacteria at high frequencies (Nelson et al., 2012; Gigliucci et al., 2018). Chen et al. (2017) found a statistically significant association between the presence of the *Enterobacteriaceae* family and patients AGE with complications, compared a control group.

The Shannon-Wiener diversity index demonstrated that from the grouping of the main phyla, there is evidence of the presence of equitability between the microorganisms present in the samples analyzed in this study, showing that these individuals had similar intestinal flora. Previous studies have indicated that a severe AGE infection causes loss of microbial diversity, which could be evidenced in the entropy score (Chen et al., 2017). Comparing samples that showed only reads for NoV and samples with reads for other viruses, we did not observe any statistically significant difference.

Previously studies of gut microbiota of healthy individuals showed the composition of the main bacterial phyla present in the intestine, represented by Firmicutes and Bacteroidetes with more than 90% of the total community, following by Actinobacteria, Proteobacteria, and Verrucomicrobia (Tap et al., 2009; Eckburg et al., 2005). Our results showed that the *Firmicutes* was most abundant bacterial phylum followed by *Proteobacteria, Actinobacteriota*, *Bacteroidota* and *Cyanobacteria,* with a taxonomic profile suggestive of a disruption of gut microbiome. NoV can alter host gut microbial communities resulting in an enhanced bacterial Firmicutes-to-Bacteroidetes (F/B) ratio, changing their role in maintaining the homeostasis in the human intestine and causing disbiosis (Tap et al., 2009).

Our results indicate that metagenomics is valuable tool for identification of the microbiota associated with NoV infection in stool samples. Besides, Bioinformatic analyzes can be directed and applied for the detection and genomic characterization of pathogens and other microorganisms of the intestinal flora, optimizing the obtaining of taxonomic profiles and provided insight into the changes occurring in the intestinal microbiota upon norovirus infection.

Studies involving the characterization of the intestinal microbiota in fecal samples from children with diarrhea are scarce, thus our data provide the first evidence of alterations in the microbiota of children with viral enteric infections in the Amazon region. However, further studies are necessary to compare the variation in taxonomic profiles with healthy children, since in this study it was only possible to analyze individuals with norovirus infection.

